# RNA Synthesis is Associated with Multiple TBP-Chromatin Binding Events

**DOI:** 10.1101/045161

**Authors:** Hussain A. Zaidi, David T. Auble, Stefan Bekiranov

## Abstract

Competition chip is an experimental method that allows transcription factor (TF) chromatin turnover dynamics to be measured across a genome. We develop and apply a physical model of TF-chromatin competitive binding using chemical reaction rate theory and derive the physical half-life or residence time for TATA-binding protein (TBP) across the yeast genome from competition ChIP data. Using our physical modeling approach where we explicitly include the induction profile of the competitor in the model, we are able to estimate yeast TBP-chromatin residence time as short as 1.3 minutes, demonstrating that competition ChIP is a relatively high temporal-resolution approach. Strikingly, we find a median value of ~5 TBP-chromatin binding events associated with the synthesis of one RNA molecule across Pol II genes, suggesting multiple rounds of pre-initiation complex assembly and disassembly before productive elongation of Pol II is achieved at most genes in the yeast genome.

---

Cellular processes including transcription are inherently dynamic. Currently, the dynamics of transcription and other molecular processes in the cell are poorly understood^1^ because of a lack of methods that measure fundamental kinetic parameters in vivo. Precise estimation of the chromatin-binding on-rates and off-rates of general transcription factors (GTFs) and other classes of transcription factors (TFs) would allow more quantitative understanding and modeling of pre-initiation complex (PIC) formation^2,3^, RNA polymerase recruitment and elongation, and transcription^4,5^. Live-cell imaging at specific multi-copy genes is capable of yielding the residence time of TF-chromatin interactions at high temporal resolution (i.e., second timescale)^6^ but in general does not allow these measurements at single-copy genes. Cross-linking kinetic (CLK) analysis is a high spatial and temporal resolution method that enables estimation of the in-vivo TF-chromatin on-rates and off-rates at single-copy loci^7,8^. Another experimental approach to assessing TF-chromatin dynamics is anchor-away (AA)^9,10^; however, only qualitative or semi-quantitative TF-chromatin dynamic information is determined from this approach^9,10^. Indeed, alternative physical-modeling approaches to calculating these kinetic parameters are needed to independently verify the estimates obtained from CLK and live-cell imaging techniques^11,12^.

Competition ChIP is another high-spatial resolution method in which the endogenous copy of a TF contains one protein tag and an alternative copy, a competitor, is transcriptionally induced with an alternative protein tag^13–15^. We developed and applied a physical modeling approach using chemical kinetic theory that directly estimates the physical half-life or residence time of TATA-binding protein (TBP)—the general transcription factor which initiates PIC formation^16^—on chromatin across the yeast genome from TBP competition ChIP data^15^. Given that the competitor TF requires 20-30 minutes for induction^14,15^, competition ChIP was generally believed to be low temporal resolution (20 minutes or greater)^7,9^. Moreover, previous analyses of competition ChIP data have estimated relative turnover rates^13–15^ and not residence times. Lickwar et al.^14^ argue that they estimated the residence time of Rap1 across the yeast genome with the shortest residence time being ~30 minutes; however, we show that their estimates, while correlated with the physical Rap1 residence time, are likely much longer than the actual physical residence time. In support of this, live cell imaging^12^, CLK^7,8^ and AA^9^ analyses reveal that TBP-chromatin interactions range from seconds to a few minutes depending on the promoter. However, the previous estimates of residence times were made at select loci using qPCR^7,9^ or represented effective averages across hundreds to thousands of promoters^12^. Consequently, this study is the first to arrive at genome-wide estimates of TF-chromatin residence times for TBP. Using our physical modeling approach, we are capable of estimating TBP-chromatin residence times as short as 1.3 minutes and as long as 53 minutes, demonstrating that competition ChIP is actually a relatively high temporal resolution method. An advantage of estimating the physical residence time as opposed to relative turnover is that comparison of physical residence times to other physical timescales including nascent RNA transcription rates inform qualitative and quantitative models of the efficiency or stochasticity of PIC formation and transcription^1^. Furthermore, physical residence times will lead to physical mathematical models of PIC assembly and transcription^2,3^ as more kinetic parameters are measured.

Comparing TBP-chromatin residence times with nascent RNA transcription rates^17^, we found that a median value of ~5 TBP binding events were associated with productive RNA synthesis across Pol II genes. Our results paint a highly dynamic, stochastic picture of pre-initiation complex formation with multiple rounds of partial assembly and disassembly before a single round of productive RNA polymerase elongation. We also compared TBP-chromatin residence times to Rap1 and nucleosome relative turnover^13–15^. Notably, these are the only other regulatory factors whose dynamics have been characterized at specific sites on a genomic scale. We found that TBP-chromatin residence time was correlated with Rap1 ^13–15^ but not nucleosome ^13–15^ turnover dynamics. Moreover, while TBP and Rap1 chromatin dynamics were poorly correlated with nascent RNA transcription rates^17^, +1 nucleosome turnover dynamics, which likely affect Pol II elongation ^18–20^, showed modest but robust positive correlation with nascent RNA transcription rates. Assessment of the role that the occupancy of over 200 transcription factors^21^ played in modulating TBP-chromatin residence times and nascent RNA transcription rates across gene promoters revealed only a subunit of TFIIE affecting TBP residence times while a number of initiation and elongation-related TFs had a relatively strong impact on nascent RNA transcription rates. Our findings point to the dynamics and occupancy of factors that regulate the late stages of transcription initiation including Pol II elongation associating more strongly with nascent RNA transcription rates than that of factors regulating early stages including PIC formation such as TBP and Rap1.

## Results

### Overview of Competition ChIP Experiment and Data Analysis

Competition ChIP (schematically represented in Fig. 1a-d) enables direct measurement of TF-chromatin turnover dynamics at binding sites across a genome (e.g., yeast genome). This is accomplished by attaching a protein tag to an endogenous TF (orange dots in Fig. 1a-d) and by expressing a competitor of that TF with a different tag (maroon dots in Fig. 1a-d). The relative occupancy of the alternatively tagged TFs are measured at binding sites across a genome using chromatin immunoprecipitation (ChIP) followed by hybridization to genomic tiling arrays (ChIP-chip) or high throughput sequencing (ChIP-seq). Quantification of the normalized ratio of induced competitor TF ChIP signal over the endogenous TF ChIP signal over time after induction of the competitor TF yields estimates of TF-chromatin turnover at any given binding site^14,15^. The induction of the competitor concentration (labeled *C_B_*) relative to the endogenous TBP concentration (labelled *C_A_*) takes ~60-70 minutes to reach steady state levels as shown by the dashed line in Fig. 1e-g.

**Figure 1.**
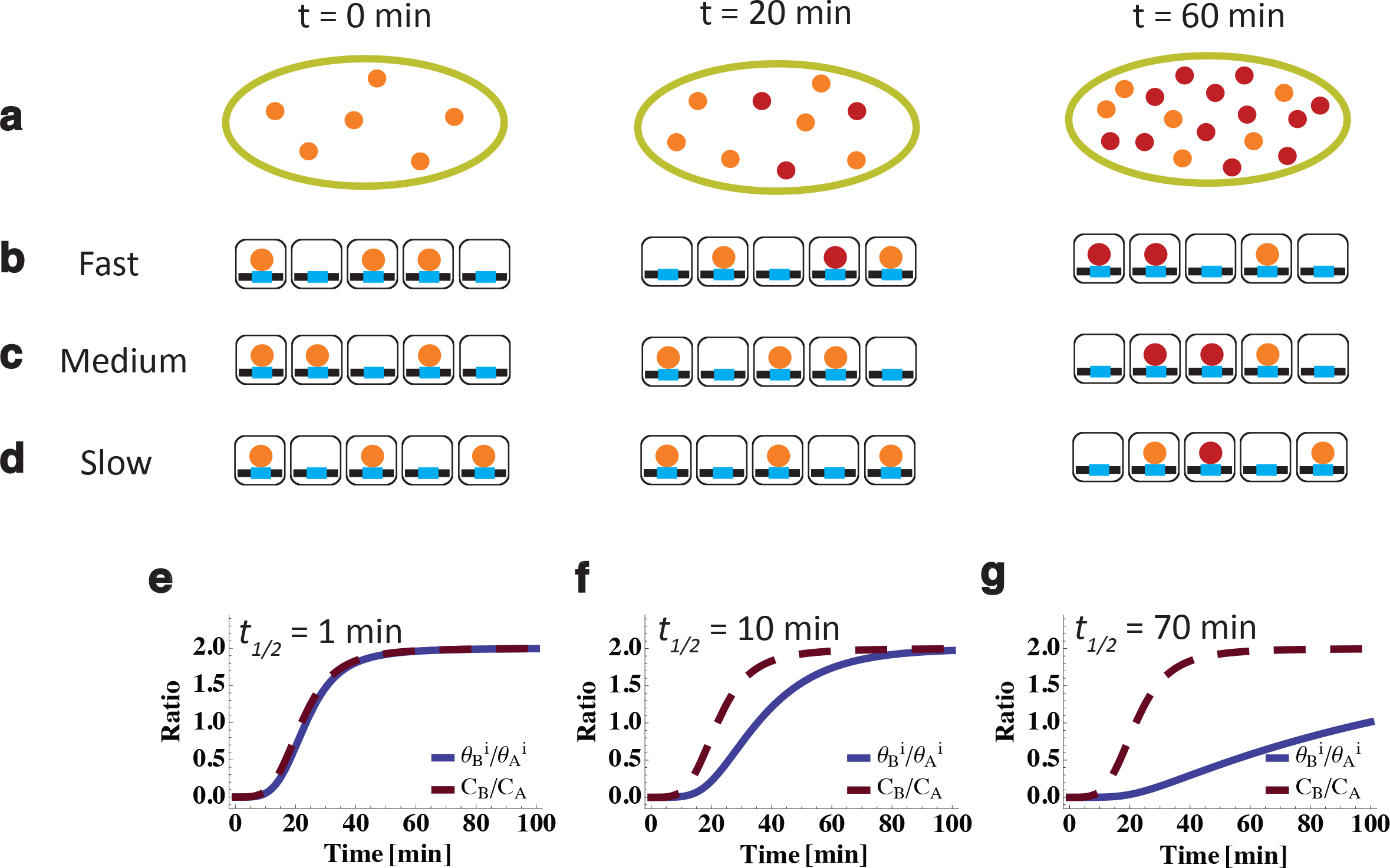
Illustration of Competition ChIP Experiment. **a-d**, In vivo induction of HA-tagged competitor TBP (maroon), in vivo stable population of Avi-tagged endogenous TBP (orange), and a depiction of fast, medium, and slow binding dynamics over induction times of 0 min, 20 min, and 60 minutes. **a,** Induced TBP concentration going from zero at 0 min to twice the endogenous TBP at 60 min of induction time, approximately following the induction in van Werven et al.^15^. The induction curve is also labeled as *C_B_/C_A_* (dashed brown curve) in **e-g**. **b-d,** The “Fast”, “Medium”, and “Slow” rows depict the binding of induced and endogenous TBP at loci with TBP residence times of less than a minute, a few minutes, and tens of minutes, respectively, for given induction times of 0 min, 20 min, and 60 min. **e-g,** Simulated in-vivo ratio of occupancy of induced to endogenous TBP with a residence time of 1 min, 10 min, and 70 min. **e,** For loci with fast dynamics, the occupancy ratio follows the induction curve closely, also depicted in **(b)** where the ratio of sites occupied by competitor to those occupied by endogenous TBP closely follows the ratio of concentrations of competitor to endogenous TBP shown in (a).**f,** The occupancy ratio lags behind the induction curve for TBP residence time of 10 min. At 20 min post-induction the ratio of occupancies is almost zero, also shown by the absence of maroon dots in the middle panel of **(c)**. Since the induction curve approaches the saturation value of 2 around 50 minutes, the ratio of occupancies starts approaching the induction curve around 60 minutes, also shown in the last panel of **(c)** where the induced TBP occupancy is twice that of the endogenous TBP. **g,** The rise and saturation of the ratio of occupancies is significantly delayed compared to the induction curve for TBP residence time of 70 min. Around 60 minutes, the ratio of induced occupancy to endogenous occupancy is ~ 0.5, also shown in the last panel of the **(d)** with induced TBP bound to one locus and endogenous TBP bound to two loci.

We applied kinetic theory to model the in vivo competitive dynamics of the induced competitor and the endogenous TBP in a competition ChIP experiment^15^ to estimate the TBP-chromatin binding on-rate (*k_a_*) and off-rate (*k_d_*) (Supplementary Text Sec. 2) at sites across the yeast genome. We found that the ratio of simulated induced over competitor occupancy versus time strongly depended on residence time (*t*_1/2_ = ln(2)/*k_d_*) and not the on-rate (Supplementary Text Sec. 3). Additionally, we observed that the simulated ratio of occupancies using the kinetic model (solid lines in Fig. 1e-g) rose and saturated (at steady state levels) at slower rates with increasing residence time, t_1/2_. Consequently, TBP-chromatin interactions with short residence times (*t*_1/2_ = 1min) yielded a simulated ratio of occupancies versus time that was mildly but noticeably displaced or shifted (i.e., minute time scale) to the right of the induction curve (Fig. 1e) while TBP-chromatin interactions with longer residence times were displaced roughly by the value of the residence time (Fig. 1f,g). Intuitively, this time-delayed response of the ratio of occupancies relative to the induction curve can be viewed as an additional delay compared to induction driven by the residence time of the TF. In fact, the simulation showed that the residence time is effectively the time it takes for the turnover to affect the ratio of occupancies in response to induction of the competitor at all times post induction, including times much shorter than that required for full induction. Importantly, this delay is noticeable as soon as the induction curve rises above 0 (i.e., noise level), which is ~10 minutes (Fig. 1e-g), and, as discussed below, enables residence times as short as ~1 to 2 minutes to be estimated.

Along with background subtraction and normalization of TBP competition ChIP data^14,15^ (Supplementary Text Sec. 1), an important data processing step includes scaling of the normalized, background subtracted ChIP ratios at steady state (i.e., *t* → ∞) as outlined in Fig.1h (also Supplementary Text Sec. 2). In order to fit a kinetic theory of competitive binding represented as a ratio of the competitor over the endogenous TF occupancies versus time, the processed data must satisfy the constraints on the ratio of occupancies at the start of induction (*t* = 0) and steady state or equilibrium (*t* → ∞). More specifically, the mathematical solution of the kinetic theory equations (Supplementary Text Sec. 2) shows that the ratio of the competitor over endogenous TF occupancies equals the ratio of competitor over endogenous TBP concentration at steady state (*t* → ∞). This is depicted in Fig. 1e-g where the ratio of simulated occupancies (solid blue lines) and the ratio of TBP competitor concentration over endogenous concentration (dashed brown line) at steady state both equal 2. Importantly, background subtraction and normalization of competition ChIP genomic tiling array or high throughput sequencing data across time points does not yield properly scaled data at steady state (as shown in Fig. 2b). There are likely multiple reasons for this discrepancy between theoretical and background-subtracted normalized ratios including differences in the affinity of the two antibodies used to tag the competitor and endogenous TF (Supplementary Text Sec. 2). Nevertheless, if a kinetic model is used to fit competition ChIP data, the data must be properly scaled to satisfy the constraints of the theory at the start of induction and at steady state—a crucial step that has not been implemented 13,14 previously.

**Figure 2.**
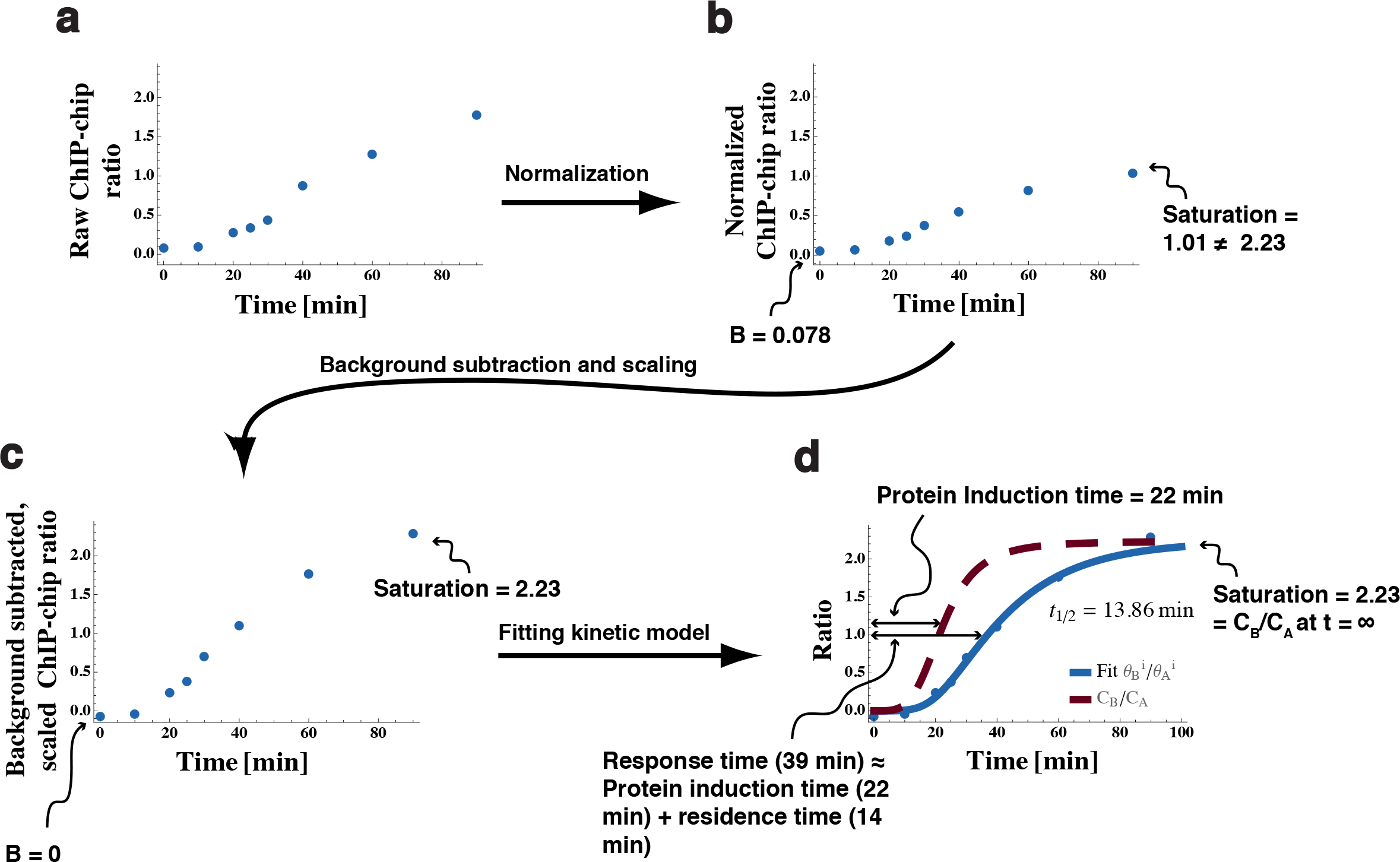
Schematic workflow of the quantitative analysis pipeline. **a-d**,A schematic representation of the data processing pipeline that takes the geometrically averaged ratio of HA and Avi proteins from van Werven et al.^15^ and outputs scaled, normalized ratios that can be fit with the in vivo kinetics model. **a,** The first step was to normalize the data for each induction time to the non-specific background to take into account potentially different experimental conditions for the time points. **b,** After normalization, a sigmoid with a constant was fitted to the data for each locus: the constant (B) gave the locus specific background value, and the amplitude gave the saturation value for the ratio data. In the figure, the locus specific background is 0.078, and the saturation value is 1.01. The expected saturation value at each locus given by the in vivo kinetic model is the ratio of the concentrations of the competitor to the endogenous TBP at long induction times (~2.23 as shown in Fig. 3a). **c,** We subtracted the background (B) from the locus data and scaled the data with a multiplicative factor such that the saturation matched the expected saturation value of 2.23, without which the data and the theory would be at odds. **d,** After data processing, the data was fitted with the in vivo kinetic model to extract residence times. A heuristic, approximate explanation of the “lag” between the induction curve and the observed occupancy ratio is that the response time (denoted by *t_0_*) as measured by fitting a sigmoid to locus data without using the kinetic model is approximately the sum of the protein induction time (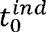) and the extracted in vivo residence time (t_1/2_) found using the kinetic model. This signifies that the residence time can be qualitatively approximated as the difference between the response time and the protein induction time.

### Background Subtraction, Normalization and Scaling of Competition ChIP-chip Data

In order to fit TBP competition ChIP two-color Agilent tiling microarray data^15^ to our kinetic model, we first normalized each dataset to non-specific background (Fig. 2b, Supplementary Fig. 1, and Supplementary Text Sec. 1). We then subtracted locus-specific background and scaled the data for TBP peaks within gene promoters to theoretically expected values at the start of induction (*t* = 0) and steady state or equilibrium (*t* → ∞) (Fig. 2c and Supplementary Text Sec. 2). The kinetic theory explicitly accounts for the time dependence of the induction of the competitor. Consequently, we fit the ratio of the induced (denoted by B) over endogenous (denoted by A) TBP concentration (*C_B_(t)/C_A_*) determined from Western blots as a function of induction time^15^ to a function that displayed critical features of the ratio: saturation as well as positive curvature (i.e., increasing slope) at low time points and negative curvature (i.e., decreasing slope) near steady state or saturation. A Hill-like sigmoid function with Hill coefficient *n* = 4 (Fig. 3a and Supplementary Eqn. 3) displays all of these properties and yielded the best fit of the ratio of concentration data over time. The fit yielded a characteristic time-scale for TBP competitor induction 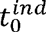 = 22 min and the steady state ratio of induced over endogenous TBP concentration (*C_B_(*t* → ∞)/C_A_* → 2.23). Not surprisingly, the normalized competition ChIP data at nearly every TBP binding site was also well approximated by an n = 4 Hill-like equation with a time-scale parameter*t*_0_ (Supplementary Eqn. 3), which quantifies the *overall* turnover response including induction and TF-turnover dynamics at every TBP peak. As we showed in our simulation of ratios of competitor over endogenous TF occupancies using kinetic theory of competitive binding (Fig. 1e-g), the resulting competition ChIP ratio (after proper normalization, background subtraction and scaling) is a response curve that is delayed compared to the induction curve (with a characteristic time-scale 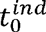) roughly by the residence time (*t*_1/2_) (i.e., crudely 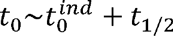). We used this Hill-like equation to background subtract and scale the data to the theoretical in vivo (denoted by superscript *i*) ratio of fractional occupancy of the competitor 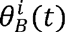 to the endogenous 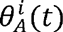 TBP, which must satisfy the boundary conditions at the start of induction (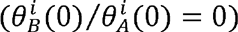) and steady state (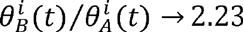as *t*→ ∞) as described above (Fig. 2 and Supplementary Text Sec. 2,4).

**Figure 3.**
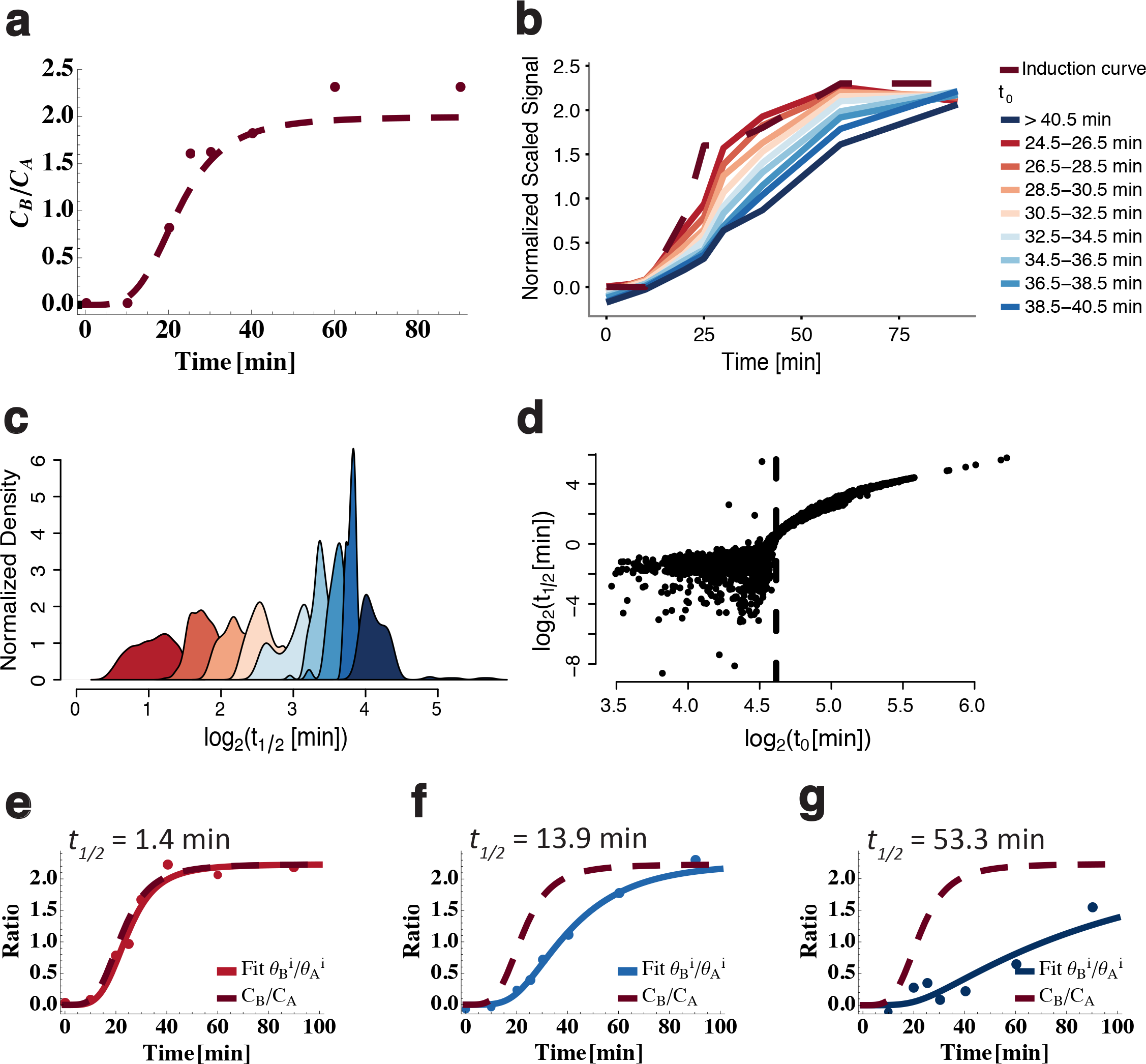
Estimation of TBP residence time from kinetic model fit to normalized, scaled competition ChIP data. **a**, Ratio of concentration of competitor TBP(*C_B_*) to the concentration of endogenous TBP (*C_A_*) taken from van Werven et al.^15^ along with a sigmoid fit to the data (dashed line). The fit gave a saturation value of 2.23 and protein induction time min (the time at which the signal reaches half the saturation value). **b,** Plot of normalized, scaled competition ChIP ratio data (competitor/endogenous) versus induction time. The dashed line shows the protein induction data from (**a**). As shown in Fig. 1, *t_0_* is an estimate of the overall turnover response time. Hence, the data stratified and averaged in bands of 2 minutes for *t_0_* ranging from 24.5 minutes to greater than 40 minutes showed a progressively slower rise as *t_0_* increased. **c,** Normalized density of TBP residence times, *t_1/2_*, obtained from data in each band (same color scheme as panel (**b**)) showing that larger leads to longer residence times as explained in Fig. 1. **d,** log_2_–log_2_ plot of TBP *t_1/2_* versus response time showing a monotonic relationship between *t_1/2_* and *t_0_* for *t_0_* > 24.5 min. For *t_0_* < 24.5 min, the noise in the data and the induction curve made *t_1/2_* estimates imprecise. As a consequence, estimates of residence times shorter than ~ 90 seconds are unreliable. **e - g,** Representative fits of our kinetic theory based model to the normalized, scaled competition ChIP ratio data and estimates of TBP *t_1/2_*, along with the fit to the protein induction data (dashed, same as **(a)**). The colors of the data and the fits correspond to the appropriate *t_0_* band shown in **(b)**. **e-g** once again highlight that the residence time extracted using the kinetic model increases as the response time increases.

### Estimation of Residence Time by Fitting the Model of Competitive Binding to Normalized, Scaled Competition ChIP Data

We then simultaneously numerically solved and fitted the in vivo kinetic equations of competitive binding between species A and B (Methods Eqns. 1 and 2, Supplementary Eqns. 9 and 10) to normalized, scaled competition ChIP data (Fig. 2 and Supplementary Text Sec. 4, 5). We (and others^13–15^) ignored the impact of cross-linking theoretically as competition ChIP data was gathered at one cross-linking time (20 min of formaldehyde cross-linking in van Werven et al.^13–15^). We showed that the resulting off-rate, *k_d_*, could be modestly biased (Supplementary Fig. 3a-d) using a generalization of the CLK framework with crosslinking to competition ChIP (Supplementary Eqns. 4-8). This framework could be used to correct the bias if data is gathered at various crosslinking times^7^. As noted by Lickwar et al.^14^, we also found that the in vivo ratio of induced over endogenous TF as a function of induction time is insensitive to the on-rate, *k_a_*, and is very sensitive to the off-rate or residence time, *t*_1/2_ = ln(2)/*k_d_* (Supplementary Text Sec. 3 and Supplementary Fig. 3e-f). Consequently, we only arrived at relatively precise values of the residence time (*t*_1/2_).

### TBP-Chromatin Residence Times Ranging from 1.3 to 53 Minutes Estimated Fromnormalized, Scaled Competition ChIP Data

Stratifying TBP-containing promoters in 2-minute bands of *t*_0_, we showed that the average normalized and scaled ratio of competitor over endogenous signals as a function of induction time progressively showed slower rise as *t*_0_ increased (i.e., moved to the right) (Fig. 3b) with corresponding residence times increasing from 1.3 to 53 minutes (Fig. 3c, Supplementary Text Sec. 6), showing that residence times could be estimated from the ratio. Indeed, given that fitting the Hill-like equation and chemical kinetic equations should yield highly correlated results, we found a smooth relationship between *t*_1/2_ and *t*_0_ (as mentioned earlier, crudely 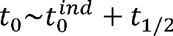) up to a point where numerically fitting the chemical kinetic equations became unstable; this point is marked by *t*_0_ < 24.5min (Fig. 3d and Supplementary Fig. 4). This numerical instability was due to the fact that for promoters with min, the separation between the normalized, scaled data and the induction curve were well within the noise of the competition ChIP data. For *t*_0_ > 24.5minmin, the normalized, scaled data yielded excellent fits to the chemical kinetic equations, the data moved progressively to the right with increasing residence time and, remarkably, allowed residence times as short as 1.3 minutes to be estimated (Fig. 3e-g, Supplementary Fig. 4e-h, and Supplementary Table 1). Importantly, the shortest residence time that could be reliably estimated was determined by the noise in the induction and competition ChIP data and not the induction time of the competitor. As soon as reliable, robust separation (i.e., beyond their relative error or noise) between the induction and competition ChIP ratio curves existed, relatively reliable residence times could be estimated.

**Figure 4.**
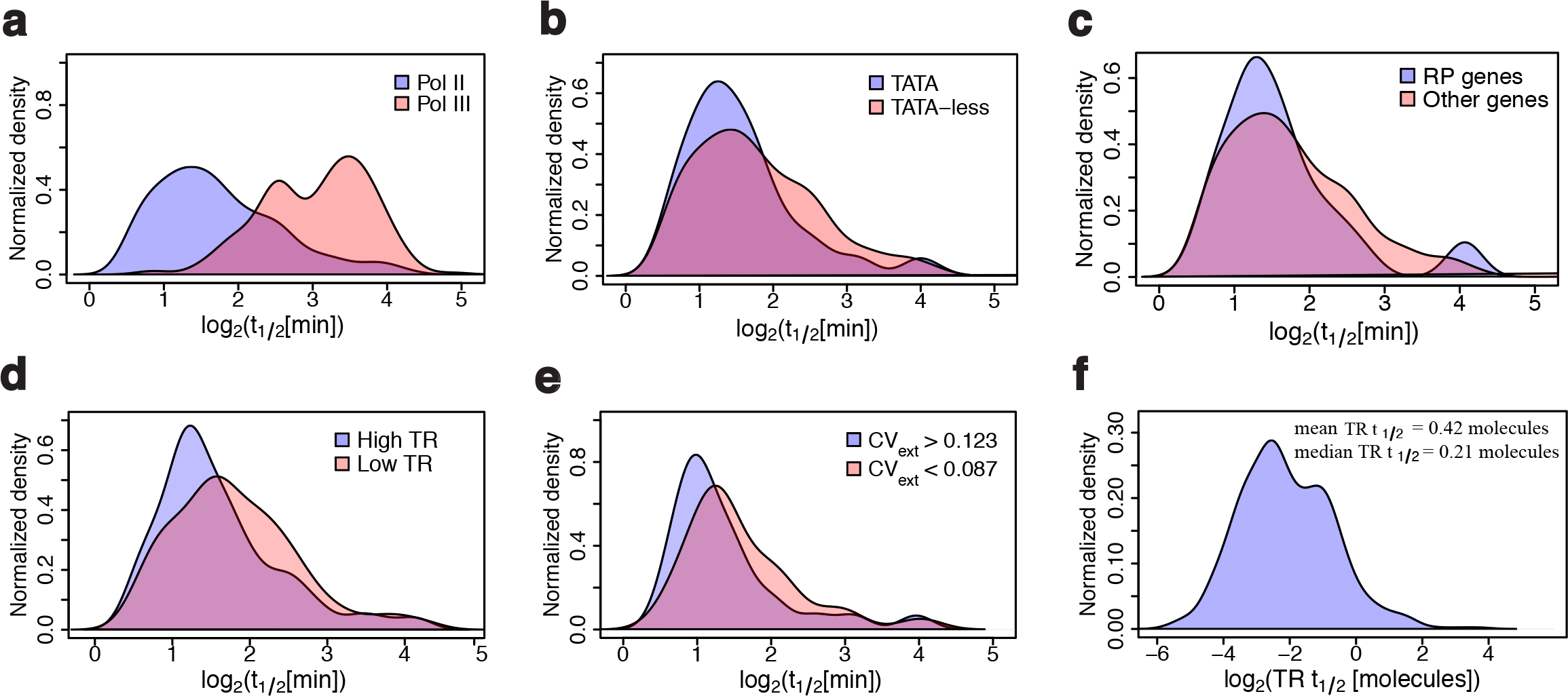
Multiple, minute-scale TBP-chromatin binding events are associated with transcription at Pol II genes. **a**, Normalized density of TBP residence time (on log_2_ scale) for Pol II and Pol II promoters which yielded a median Pol II TBP residence time (*t_1/2_*) of 3 min and median for Pol III genes of 9 min. The difference between the two distributions is significant with a Kolmogorov-Smirnoff (KS) p-value =2.2e-16. **b,** Normalized TBP *t_1/2_* (on log_2_ scale) density for TATA-containing versus TATA-less promoters. TATA-containing promoters have over all shorter residence times than TATA-less promoters (KS p-value = 0.0075). **c,** Ribosomal protein (RP) genes have marginally shorter TBP residence times compared to non-RP genes (median RP*t_1/2_* = 1.4 min and median non-RP *t_1/2_* = 1.6 min; KS p-value=0.25). **d,** Promoters in the highest quartile of transcription rate (TR) tend to have shorter TBP *t_1/2_* than promoters in the lowest quartile (KS p-value = 0.005). **e,** Promoters with higher extrinsic transcriptional noise (*CV_ext_*)^30^ have lower TBP residence time (KS p-value = 0.048). **f,** Normalized density of transcription efficiency (defined as the transcription rate multiplied by residence time, *TR t_1/2_*) showing that the median transcriptional efficiency is 0.21 molecules. In other words, for a representative Pol II promoter, ~5 TBP turnovers are required before a single molecule of RNA is successfully transcribed (inverse of transcriptional efficiency).

### Multiple TBP-Chromatin Binding Events are Associated with Synthesis of One Nascent RNA Molecule at Pol II Genes

Earlier estimates of relative TBP turnover, *r* for 602 Pol II and 264 Pol III genes were obtained using linear regression to a subset of the data (i.e., 10, 20, 25 and 30 min time points)^15^. Because a physical model of competitive binding rooted in reaction-rate theory naturally follows the profiles of the normalized and scaled data as a function of induction time (as opposed to a linear fit), we were able to apply stringent noise criteria on the residuals of each fit (Supplemental Text Sec. 8) and reliably estimate TBP residence times for 794 Pol II and 205 Pol III genes (Supplementary Table 1). While *r* and our estimates of *k_d_* are correlated (Supplementary Fig. 5a), *r* is also strongly correlated with the *t* = 0 ratio of induced over competitor ChIP signals (Supplementary Fig. 5), which suggests insufficient background subtraction influencing the estimates of *r*. Nevertheless, in agreement with estimates of *r* made by van Werven et al.^15^ as well as the competition ChIP and AA results of Grimaldi et al.^9^, we found that TBP residence times were notably shorter for Pol II compared to Pol III genes (Fig. 4a) and to a lesser extent for TATA compared to TATA-less genes^22^ (Fig. 4b). In contrast to van Werven et al.^15^ but consistent with Grimaldi et al.^9^, we found no significant differences between TBP residence times comparing SAGA containing and SAGA free genes (Supplementary Fig. 6d) or TFIID-containing and TFIID-free genes (Supplementary Fig. 6g). Given that Pol III genes tend to be higher expressed^23^ and have longer TBP residence times than Pol II genes, we were surprised to find marginally shorter TBP residence times at highly expressed ribosomal protein (RP) genes compared to other genes (Fig. 4c, Supplementary Fig. 6j). This finding was consistent with higher nascent RNA transcription rates (TRs)^17^ for shorter TBP residence times at Pol II genes (Fig. 4d). Shorter residence times were also associated with higher levels of extrinsic transcriptional noise^24^ (Fig. 4e) consistent with recent findings^25^. With estimates of TR and TBP *t*_1/2_, we defined *transcriptional efficiency*, which is the product of the transcription rate and TBP residence time (*TRt*_1/2_) and represents the number of TBP residence times or binding events associated with productive elongation of Pol II and transcription. Strikingly, we found low transcriptional efficiencies for Pol II genes (Fig. 4f). The median *TRt*_1/2_ across Pol II promoters was 0.2, or ~5 TBP binding events for productive RNA synthesis to proceed (Fig. 4f). This is consistent with an upper limit for this value for most Pol II genes (i.e., *TRt*_1/2_ ≤ 1) determined by the likely TBP-chromatin residence time from AA experiments and characteristic values of transcription rate across the yeast genome^9^. These findings are consistent with rapid, highly stochastic TPB/PIC dynamics at Pol II genes with multiple rounds of assembly and disassembly before productive Pol II elongation. While we don’t have nascent RNA data for Pol III genes, these genes tend to be much higher expressed than Pol II genes; yet TBP residence times tended to be ~10 minutes (Fig. 4a) suggesting much more stable PIC formation^26^ and function for Pol III genes.

**Figure 5.**
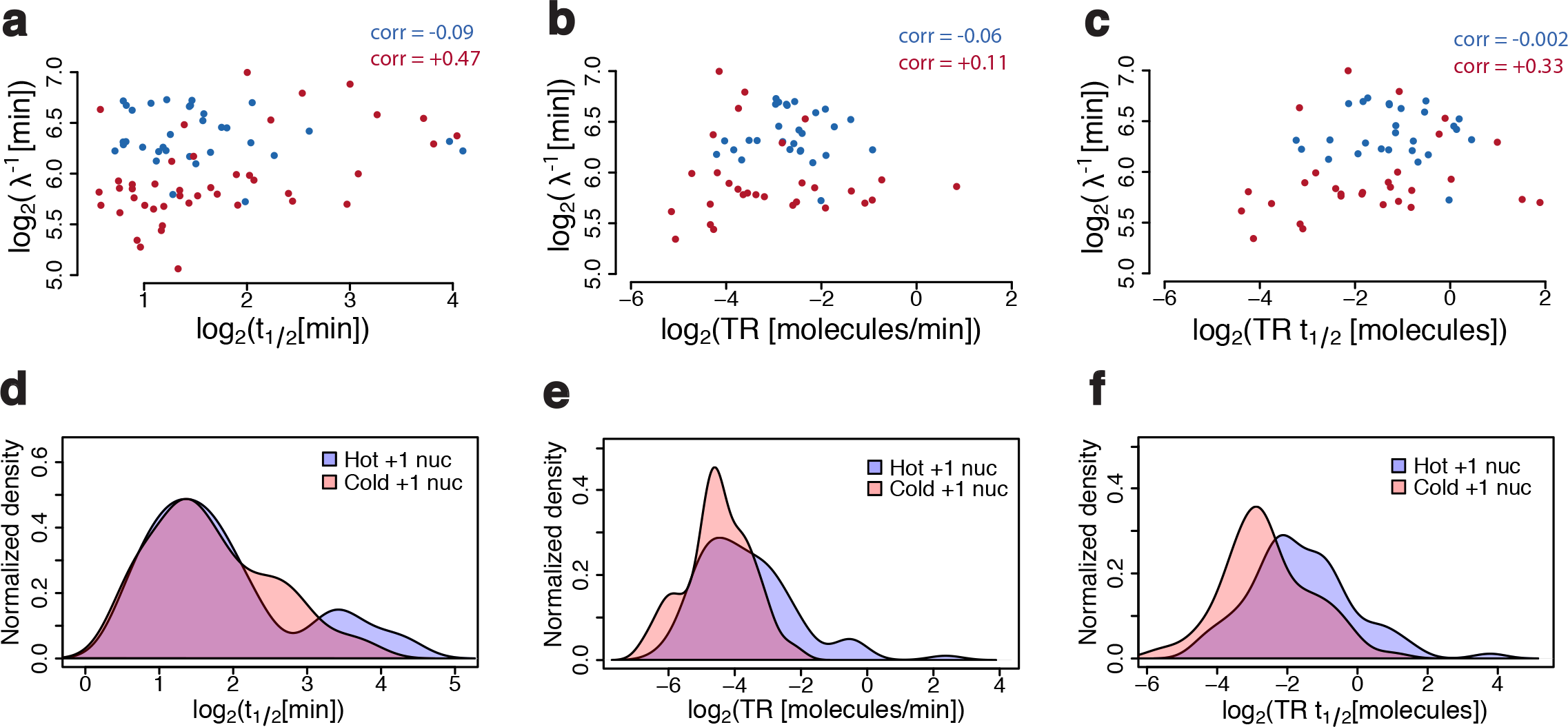
TBP dynamics are correlated with Rap1 but not +1 nucleosome dynamics. **a-c**, log_2_–log_2_ scatterplot of Rap1 relative residence time (*λ*^-1^) versus **(a)** TBP residence time (*t_1/2_*), **(b)** transcription rate (*TR*), and **(c)** transcription efficiency (*TRt_1/2_*) for Ribosomal protein (RP) genes in blue and non-RP genes in red. Rap1 *λ*^-1^ correlated well with TBP *t_1/2_* and *TRt_1/2_* at non-RP genes, but not at RP genes. *λ*^-1^ was mildly correlated with *TR* at RP genes. **d-f,** Normalized density of **(d)** TBP residence time (*t_1/2_*), **(e)** transcription rate (*TR*), and **(f)** transcription efficiency (*TRt_1/2_*) at genes containing hot and cold +1 nucleosomes. Hot nucleosomes were in the top quartile of nucleosome turnover and cold were in the bottom quartile (see Supplementary Text Sec. 15). There is no difference in TBP *t_1/2_* between hot and cold nucleosomes (KS p-value=0.50) (**d**), but hot nucleosomes tend to have higher *TR* (KS p-value=0.007) (**e**) and higher *TRt_1/2_* (KS p-value=1.3e-7) (**f**).

**Figure 6.**
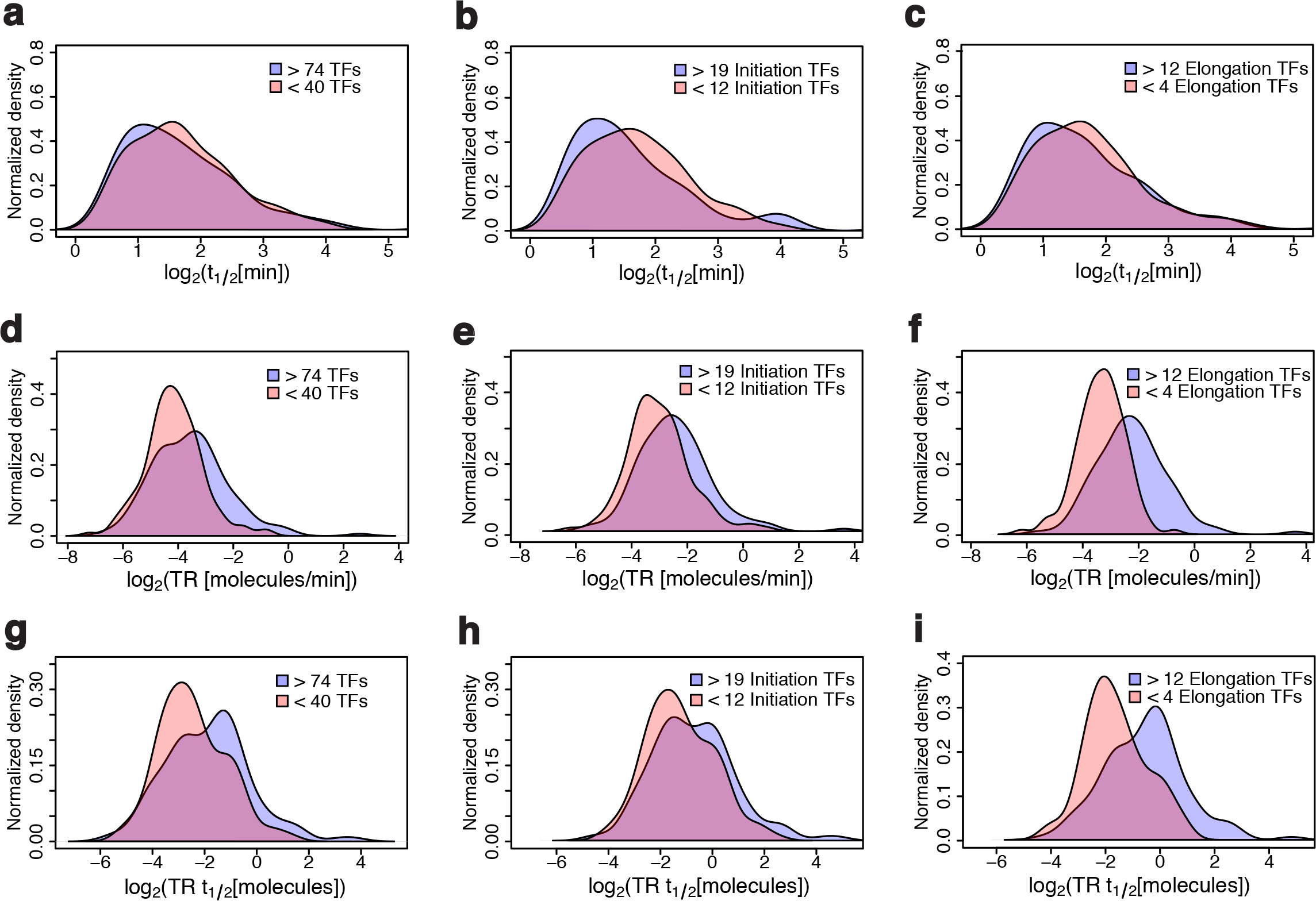
High numbers of elongation factors at Pol II promoters are associated with higher transcription rates and efficiencies. **a-c,** Normalized density of TBP residence time (*t_1/2_*) on log_2_ scale for genes with the upper quartile numbers of bound transcription factors (TFs) and genes with the lower quartile numbers of bound TFs (out of 202 mapped TFs in Venters et. al^21^) showing that *t_1/2_* is not modulated by **(a)** the number of total TFs, **(b)** initiation TFs or elongation TFs (**c**). The elongation and initiation TFs were annotated as in Venters et. al^8^. **d-f,** Normalized density of transcription rate (*TR*) on the log_2_ scale for genes with the upper quartile numbers of bound TFs and genes with the lower quartile numbers of bound TFs showing that *TR* is modulated by **(d)** the number of total TFs (KS p-value=8.6e-5), **(e)** initiation TFs (KS p-value=9.8e-4), and **(f)** elongation TFs (KS p-value=3.14e-11). **g-i,** Normalized density of transcription efficiency (*TRt_1/2_*) on log_2_ scale for genes with the upper quartile numbers of bound TFs and genes with the lower quartile numbers of bound TFs showing that *TRt_1/2_* is significantly modulated by **(g)** the number of overall TFs (KS p-value=2.4e-4), **(h)** initiation TFs (KS p-value=0.05), and **(i)** elongation TFs (KS p-value=8.5e-8) (**i**).

### TBP-Chromatin Residence Time is Correlated with Relative Rapl Residence Time but not with +1 Nucleosome Residence Time or Nascent RNA Transcription Rate

To gain further insights into the upstream regulation and/or downstream impact of TBP-chromatin binding dynamics especially on regulation of gene expression, we compared TBP residence times (*t*_1/2_) to the only other regulatory factors whose dynamics have been characterized on a genomic scale (in yeast): previously derived Rapl^14^ and nucleosome^13^ relative turnover rates (*λ*) and, their inverse turnover rates (*λ*^-1^) or relative residence times. Notably, we showed that the relative turnover (*λ*), derived using a Poisson statistical turnover model^13,14^, equals the off-rate (*k_d_*) plus a time-dependent function (Supplementary Eqn. 28, Supplementary Fig. 7) and can be moderately biased. More importantly, the relative turnover rates are excessively biased because normalized ChIP ratios were not scaled to ratios of fractional occupancies before model fitting^14^ as described above (Fig. 2, Supplementary Text Sec. 14 and Supplementary Fig. 8). In other words, fitting a model of the ratio of occupancies to un-scaled data (Fig. 2b) as opposed to properly scaled data (Fig. 2c) yields significantly biased (i.e., 30-fold or greater) estimates of Rap1 residence time (Supplementary Text Sec. 14 and Supplementary Fig. 8). That said, we found TBP residence time (*t*_1/2_) was correlated with Rap1 relative residence time (*λ*^-1^) at non-RP Pol II genes but not at RP Pol II genes (Fig. 5a). TBP residence time showed weak negative correlation with Pol II transcription rate (corr = −0.11; Supplementary Fig. 9a). Rap1 relative residence time (*λ*^-1^) showed slight positive correlation (Fig. 5b) with transcription rate at non-RP genes, while transcriptional efficiency was modestly correlated with Rap1 relative residence time at non-RP Pol II genes (Fig. 5c). Interestingly, the majority of the sites for which Rap1 relative residence times have been determined (ranging from 30-150 min) exhibit highly dynamic TBP (*t*_1/2_ < 1.4 min or *t*_0_ < 24.5 min; Supplementary Fig. 9b). This further supports our findings that Rap1 relative residence times^14^ are 20 to 30 fold higher (or more) than, but likely correlated with, actual Rap1 residence times^14^ (Supplementary Text Sec. 14). While +1 nucleosome dynamics were poorly correlated with TBP residence time (Fig. 5d, Supplementary Fig. 9c,d), they were positively correlated with transcription rate (Fig. 5e, Supplementary Fig. 9e) and efficiency (Fig. 5f, Supplementary Fig. 9f). These results suggest that while the dynamics and not merely the presence (Supplementary Fig. 6m) of transcription factors like Rap1 regulate TBP/PIC dynamics, TBP and Rap1 recruitment and dynamics are not the rate-limiting step in transcription at Pol II genes. Conversely, the dynamics of factors that play a role in regulating elongation including +1 nucleosome turnover^18–20^ may play more critical roles in determining the transcription rate and efficiency.

**Figure 7.**
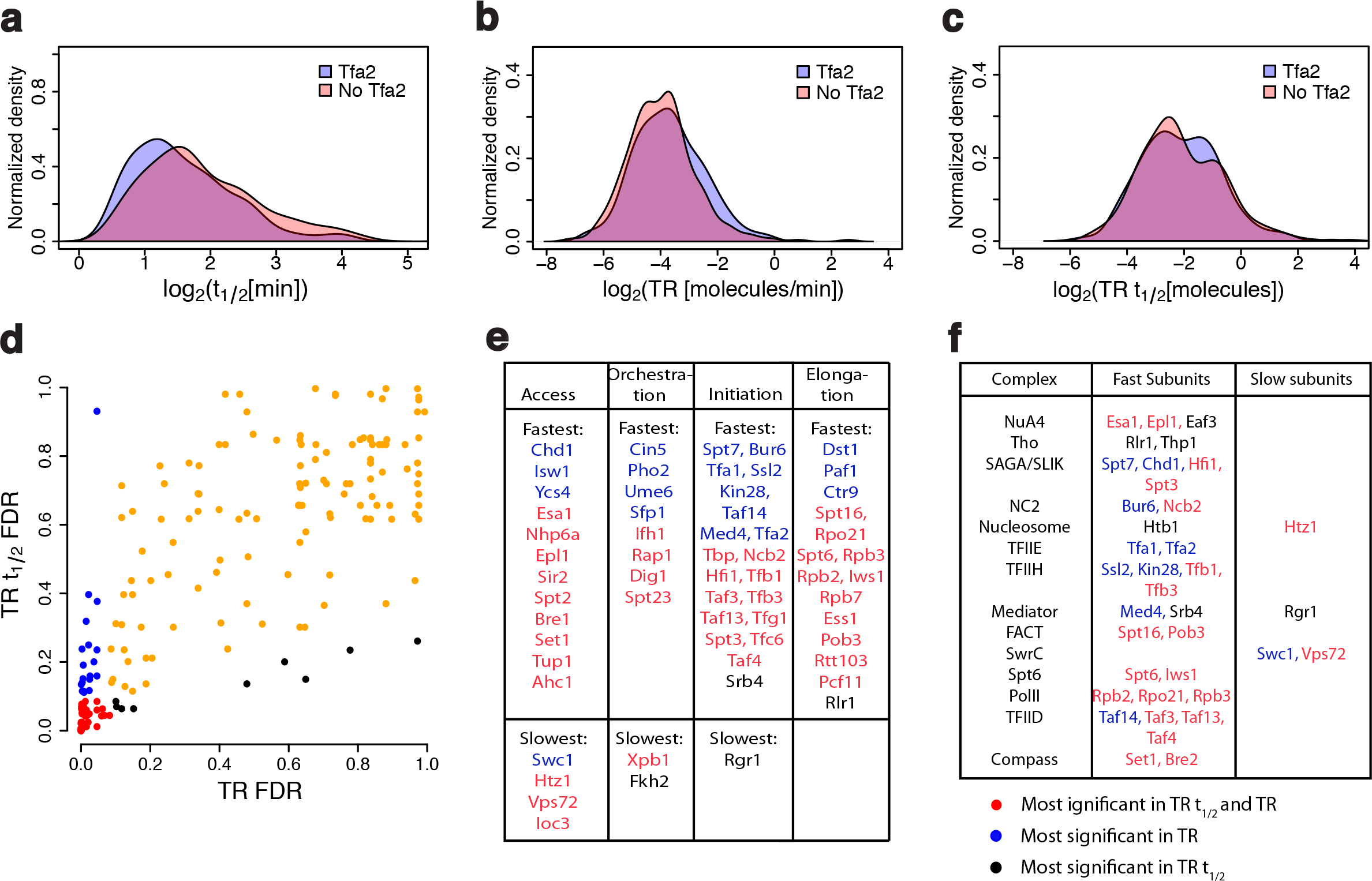
Presence of TFIIE is associated with lower TBP residence times. **a-c,** The presence of Tfa2 yielded **(a)** lower TBP residence time *(t*_1/2_) (KS p-value=5.4e-3) and **(b)** transcription rate (*TR*) with KS p-value=0.028, while **(c)** the transcription efficiency (*TRt*_1/2_) was not affected by Tfa2. **d,** Significance of the genome-wide change in TR (x-axis) and *TRt*_1/2_ (y-axis) due to the presence or absence of each TF (each dot) of 202 mapped TFs^21^. The false discovery rate (FDR) was calculated by performing a permutation t-test between the loci where a TF was bound and the loci that lacked that TF, after which multiple hypothesis correction (Benjameni-Hochberg correction) was applied. Low FDRs were used to identify TFs that are likely to affect the transcription rate only (blue dots), the transcription efficiency only (black dots) or both (red dots). **e,** Ranked list of TFs plotted in **(d)** (categorized according to access, orchestration, initiation and elongation^21^) with the highest (“Fastest”) or lowest (“Slowest”) *TR* (blue), *TRt*_1/2_ (black), or both *TR* and *TRt*_1/2_ (red).**f,** Select multi-protein complexes from the list in (**e**) highlighting the role of each complex in increasing (“Fast Subunits”) or decreasing (“Slow Subunits”) *TR* and/or *TRt*_1/2_.

### Occupancy of Multiple Elongation and Initiation Complexes at Promoters Tends to Increase Transcription Efficiency and Rate but does not Affect TBP-Chromatin Residence Time

To further assess the hypothesis that transcription factors associated with elongation as opposed to PIC and Pol II recruitment or initiation are the rate-limiting step in transcription, we tested the effect that the presence or absence of 202 transcription factors mapped to the yeast genome^21^ had on TBP residence time, transcription rate and transcription efficiency. We subdivided loci for which we had estimates of TBP residence time into quartiles of the number of transcription, initiation, and elongation factors bound based on the classification by Venters et al.^21^. As expected, the presence of greater numbers of transcription, initiation and elongation factors at promoters had no significant impact on TBP residence times (Fig. 6a-c) but yielded higher transcription rates (Fig. 6d-f) and efficiencies (Fig. 6g-i). Strikingly, the presence of more elongation factors had a much greater impact on both transcription rate (Fig. 6f) and efficiency (Fig. 6i) compared to that of initiation factors (Fig. 6e,h), consistent with our hypothesis.

For each of the 202 factors, we also conducted permutation tests to estimate the significance of differences of TBP residence times, transcription rate and efficiency at sites with the factor present compared to sites with that factor absent. We only found one factor, Tfa2 (a TFIIE subunit), whose presence yielded statistically shorter TBP residence times compared to its absence (Fig. 7a). Given that TFIIE (together with TFIIH) recruitment leads to a complete PIC, which then requires ATP for formation of the transcription bubble and subsequent Pol II elongation^27^, higher occupancy of TFIIE could lead to more rapid rates of Pol II elongation and PIC disassembly. This could explain shorter TBP residence times for promoters with higher levels of TFIIE. In partial agreement with this, presence of Tfa2 at promoters modestly increased transcription rate (Fig. 7b) but had no significant effect on efficiency (Fig. 7c). In contrast, we found that 46% and 50% of all the initiation and elongation factors mapped, respectively, significantly modulated transcription rate and efficiency (Fig. 7d,e and Supplementary Text Sec. 16). Not surprisingly, many of these factors were members of initiation and elongation complexes whose enrichment at promoters lead to both increased transcription rate and efficiency (Fig. 7f). These included subunits of the PIC assembly complexes TFIID, TFIIF, TFIIH and Pol II, while enrichment of TFIIE at promoters displayed increased transcription rate only (Fig. 7e,f). In addition, presence of multiple factors in complexes associated with elongation as well as initiation at promoters including Spt6, FACT and Mediator showed higher transcription efficiency and rate (Fig. 7e,f).

Components of complexes in the “access” class—those that regulate histones and chromatin^21^—associated with establishing active states of chromatin including monoubiquitination of H2BK123 (Bre1 and Htb1), trimethylation of H3K4 (COMPASS), acetylated histone tails (SAGA, NuA4 and Ada) and chromatin remodeling (Isw1 and Nhp6a) were enriched at genes with higher transcription efficiency and rate (Fig. 7e,f). In contrast, we also found access factors associated with chromatin and transcriptional silencing including a histone deacetylase (Sir2), and proteins that directly interact with histones (Tup1 and Spt2) at genes with higher transcription rates and efficiencies (Fig. 7e). Moreover, we found the presence of the histone variant Htz1 and components of the SWR1 complex, which exchanges H2A.Z (Htz1) for H2A in chromatin, at promoters were associated with lower transcription rates and efficiencies (Fig. 7e,f). While this result is consistent with genomic studies that show an inverse correlation between Htz1 and gene expression levels^18^, deletion of Htz1 reduces the rate of Pol II elongation 24% at the *GAL10p-VPS13* gene^28^. Finally, in the “orchestration” class—sequence specific activators and repressors^21^—we found that the presence of factors (Sfp1, Ifh1 and Rap1) that are associated with activating ribosomal protein and biosynthesis genes resulted in higher transcription rates and efficiencies while a factor that activates phosphatase metabolism (Pho2) only increased transcription rates (Fig. 7e). Interestingly, factors that recruit the repressive Tup1-Cyc8 complex (Cin5 and Skn7) along with Tup1 were associated either only with increased transcription rate or both increased transcription rate and efficiency (Fig. 7e). We note that an important caveat in this analysis, which may explain the enrichment of a few transcriptional repressors at genes with higher transcription rates and efficiencies, was that the transcription rates and efficiencies were calculated for yeast in galactose^15,17^ while the 202 TFs were mapped in yeast in YPD^21^.

## Discussion

We developed and applied a physical model of competitive binding using chemical kinetic theory to TBP competition ChIP-chip data and derived TBP-chromatin residence times genome-wide in yeast. While competition ChIP was believed to be a low time resolution approach given the 20-30 minutes that it takes to induce the competitor to a concentration approaching steady state levels, we found that we could reliably extract residence times as short as 1.3 minutes. Consistent with live cell imaging^12^, CLK^7^, and AA^9^ results, many promoters displayed highly dynamic TBP with residence times less than 1.3 minutes, which could not be accurately estimated (Supplementary Text Sec. 8).

Comparison of reliable TBP-chromatin residence times, which ranged from 1.3 minutes to 53 minutes, across different promoter classes revealed highly dynamic TBP at Pol II genes and less so at Pol III genes similar to previous studies using competition ChIP^9,15^ and AA^9^. In contrast to the findings of van Werven et al.^15^, we did not find that the occupancy of SAGA or TFIID at promoters significantly modulated TBP residence time, consistent with an independent study applying both competition ChIP and AA at select loci^9^. We did find a significant but modest decrease in TBP residence time at TATA containing compared to TATA-less promoters in agreement with van Werven et al.^15^. We also found that the TBP relative turnover parameter (*r*) derived by van Werven et. al.^15^ was biased by the HA/Avi ratio at the start of induction with higher HA/Avi ratios yielding lower relative turnover values (Supplementary Fig. 5b). This could explain the discrepancy between our results and that of van Werven et al.^15^.

We also assessed the effect that the occupancies of 202 mapped TFs^21^ had on TBP residence time, transcription rate and transcription efficiency. We only found that the presence of one factor, Tfa2 (a subunit of TFIIE), significantly modulated TBP residence time: the presence of Tfa2 at promoters by ChIP-chip analysis^21^ was associated with shorter TBP residence times. Notably, the presence of the other TFIIE subunit, Tfa1, did not have an effect on TBP residence time. Based on the analyses of Venters et al.^21^, Tfa1 was present at most promoters (4350 sites)—nearly twice as many as Tfa2 (2605 sites). Thus, Tfa2 site enrichment may be a surrogate for overall TFIIE enrichment at promoters. Conversely, we found that the presence of a number of factors classified as “access”, “orchestration”, “initiation” and “elongation” by Venters et al.^21^ significantly affected—mostly increasing—transcription rate and efficiency, with the presence of multiple factors annotated as “elongation” associated with notably higher transcription rates and efficiencies than those annotated as “initiation” (Fig. 6e,f,h,i). We note that an important caveat to these conclusions is that while these annotations are useful and may indicate a predominant role for a number of these factors, many, for example FACT, play multiple roles including both “initiation” and “elongation”^18^. Another important caveat of this analysis is that our residence time, transcription rate and transcription efficiency estimates were generated from yeast in galactose while Venters et al.^21^ mapped these 202 TFs from yeast in YPD. While there are likely many genes whose TBP, PIC and transcription dynamics are not altered by the differences in media, there are some whose regulatory factor and transcription dynamics are significantly altered. This may explain the presence of a small number of repressors (Sir2, Tup1, and Spt2) from both the “access” and “orchestration” class significantly increasing transcription rate and efficiency in our study. Indeed, transcription rates and efficiencies might increase for yeast in galactose compared to YPD at genes where these factors are present at their promoters in YPD and absent in galactose.

While the presence or absence of Rap1 did not have a significant effect on TBP residence time, Rap1 relative residence time^14^ (i.e., inverse turnover rate) was correlated with TBP residence time. This suggests the possibility of a number of unknown dynamic relationships between regulatory factors that require characterization of the dynamics as opposed to static snapshots of relative occupancy determined by ChIP-seq or ChIP-chip. We also found that Rap1 residence times were likely much shorter than previously reported^14^ and likely similar to TBP residence times, consistent with findings that Rap1 activates transcription by interacting directly with the TBP-containing TFIID complex^9,29^. Neither Rap1 relative residence time nor TBP residence time was correlated with nascent RNA transcription rate or +1 nucleosome inverse turnover. However, +1 nucleosome turnover rate was positively correlated with transcription rate and efficiency. Moreover, in agreement with the conclusion of Grimaldi et al.^9^ that at least one round of PIC assembly is required for Pol II recruitment and elongation at most Pol II genes, we found a median value of ~5 TBP residence times are associated with one productive elongation of Pol II across Pol II genes (i.e., median transcription efficiency, *TRt*_1/2_, of 0.2 molecules) suggesting multiple PIC assembly and disassembly events before synthesis of one RNA molecular at Pol II genes. Taken together, these findings suggest increased dynamic coupling of TFs and GTFs at similar stages of PIC assembly, Pol II recruitment and elongation, and transcription; the dynamics of factors that are more involved in the early stages of transcription initiation including Pol II elongation (e.g., +1 nucleosome^18–20^) are likely better dynamically correlated with transcription rate. Our study highlights the importance of developing methods that estimate TF-chromatin dynamic parameters including residence time and the resulting insights that can be gained into the inherently dynamic and stochastic process of transcription. These approaches and measurements should ultimately allow the stochastic processes of pre-initiation complex formation, Pol II recruitment and elongation, and transcription to be characterized quantitatively.

## Supplementary information

accompanies this paper at www.nature.com/naturecommunications.

## Acknowledgements

We would like to thank members of the Bekiranov lab, particularly Brian Capaldo, and the Auble lab, especially Elizabeth Hoffman, for helpful suggestions. This research was supported by NIH grant R21 GM110380 (awarded to S.B and D.T.A) and NIH grant R01 GM55763 (awarded to D.T.A).

## Author Contributions

S.B. conceived the approach and guided method development and data analysis. H.A.Z. developed and applied the methods and performed the data analysis. D.T.A. made critical analysis suggestions. H.A.Z., D.T.A. and S.B. interpreted the results and wrote the paper.

## Author Information

The authors declare no competing financial interests. Correspondence and requests for materials, software and information should be addressed to S.B. or H.A.Z.

## Methods

### Background Subtraction, Normalization and Scaling of Competition ChIP Data

The raw data generated by van Werven et al.^15^ (ArrayExpress E-M-TAB-58) reported the optical signal intensity for induced (S_HA_) and endogenous (S_Avi_) TBP concentrations hybridized on an Agilent whole-genome microarray. S_HA_ and S_Avi_ were replicated by swapping Cy_3_ and Cy_5_ dyes to take into account dye-specific variations in the intensity of the optical signal. We geometrically averaged the two dye-swapped ratios (call it *R_m_*), as described in Supplemental Text Sec. 1. Non-specific background probes were identified by fitting a normal curve to the right edge of the *t* = 0 minute log_2_(*R_m_*) data as shown in Supplementary Figure 1. We selected signal probes in the tail of the normal fit to the nonspecific background with a false discovery rate (FDR) of 0.05 or less in the *t* = 0 minute data. values were normalized (denoted by 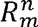) across time points, *t*, by dividing *R_m_* by the background mean obtained from the normal fit to the background probes (Supplementary Figure 1). To quantify the induction of HA over time, we fitted a Hill-like sigmoid curve with *n*=4 to the ratio of the concentration of HA to Avi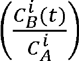 where A and B denote Avi and HA, respectively. The fit gave an induction time (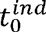) of 22 minutes and the saturation value of HA/Avi concentration ratio of 2.23 (Supplemental Eqn. 3, and Fig. 3a). We theoretically related the empirical values of 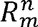 for the signal probes in our data to the ratio of the in vivo fractional occupancy of HA (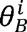) and Avi 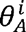 as 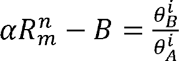, where *B* is the locus-specific differential background between HA and Avi at *t* = 0 minutes and denotes a scale factor which effectively quantifies the ratio of the antibody affinities for HA and Avi (Supplemental Text Sec. 2). To determine *α* and *B* at every TBP peak, a Hill-like sigmoid curve (with) with the added term *B* was fitted to 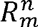 for each peak (Supplemental Eqn. 24). *B* was subtracted from 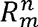 and *α* was determined as the ratio of the asymptotic in vivo concentration ratio of HA/Avi (2.23) over the asymptotic 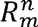 value. Hence, after scaling and background subtraction, 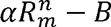 satisfied the two boundary conditions: 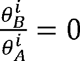 for *t* = 0, and 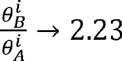 as *t* → ∞, as required by the kinetic model of in vivo competitive binding.

### Estimation of Residence Time by Fitting a Chemical Kinetic Theory Model of Competitive Binding to Normalized, Scaled Competition ChIP Data

The model for in vivo competitive binding dynamics between endogenous Avi (subscript A) and competitor HA (subscript B) TBP is described by mass-action differential equations linear in the TBP-chromatin association rate (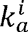) and dissociation rate (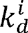):

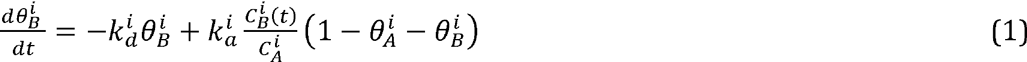

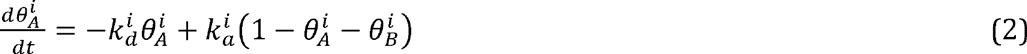

We have assumed that the association and dissociation rates for endogenous and competitor TBP are the same, superscript *i* denotes “in vivo”, and we have absorbed the experimentally undetermined endogenous concentration (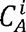) into 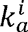, such that 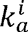 and 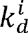 have units of inverse minutes (Supplementary Text Sec. 2). Equations (1) and (2) could not be solved analytically due to the time dependence of 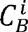, but a solution could be derived assuming that the induction of HA was instantaneous, i.e. 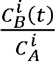 = 0 for *t* < 0 and 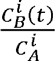 = *constant* for *t* ≥ 0. Inserting the actual time dependent 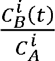 in the ideal induction solution gave an approximate solution to Equations (1) and (2) (Supplemental Eqns. 19 and 20).

We fitted the approximate solution of ideal induction to the normalized, scaled ratio data by developing a procedure for estimating the starting values for nonlinear regression to proceed (Supplementary Text Sec. 5.1). The algorithm was implemented in Mathematica and the NonlinearModelFit function with appropriate starting values was used for fitting. The ratio data is almost insensitive to 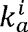 (Supplementary Text Sec. 3), and hence, we could reliably only extract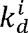. The ideal solution introduced a bias in our estimate of 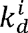, which we fixed using a pre-generated look-up table (Supplementary Text Sec. 5.2, Supplementary Fig. 2). Finally, we used our bias-corrected estimates from the look-up table as the starting point for a numerical one-dimensional Newton’s method fit of Equations (1) and (2) to find the minimum of the fit residual and extracted 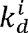 (Supplementary Text Sec. 5.3). To calculate the derivative of the fit residual required at each iteration of Newton’s method, we numerically solved the in vivo differential equations using NDSolve in Mathematica. Exceptions to the fitting procedure where we had to change the starting estimate of 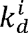 or the step size for Newton’s method are noted in Supplementary Text Sec. 5.

### Statistical Analyses of Residence Time, Transcription Rate and Transcription Efficiency Data

Throughout the main text and the supplement, quoted correlations are Spearman correlation coefficients unless otherwise stated. Kolmogorov-Smirnoff (KS) test was conducted in R using the ks.test function to determine the p-values reported in Figs. 4, 5 and 6 of the main text. For Fig. 7d of the main text, permutation test (which is useful in particular when the test statistic does not follow a normal distribution) was used to calculate the false discovery rate (FDR) for *t*_1/2_,*TR*, and *TRt*_1/2_. In other words, loci across the genome were partitioned into two sets for each transcription factor: those that showed a significant enrichment of the transcription factor above the background as determined by Venters et al.^21^ and those that did not. These two sets were used to conduct permutation test for *t*_1/2_, *TR*, or *TRt*_1/2_ test statistics using permTS in the perm library in R, which gave the mean difference of the test statistic between the two sets along with the p-value for the mean difference. The p-value was adjusted using the Benjamini-Hochberg correction^30^ using the p.adjust function in R to derive FDR estimates. In Fig. 7d the FDR for *TRt*_1/2_ was plotted against the FDR for *TR*, and transcription factors were listed in descending order of *TRt*_1/2_ mean differences. The blue dots (representing TFs that affect TR more significantly than *TRt*_1/2_) were chosen with a TR FDR < 0.06 and *TRt*_1/2_ FDR > 0.1. Red dots (representing TFs that were significant in permutation tests for both TR and *TRt*_1/2_) were chosen with TR FDR < 0.1 and *TRt*_1/2_ FDR < 0.1. Finally, black dots (representing TFs that potentially affect *TRt*_1/2_ more than TR) were chosen with TR FDR > 0.1 and *TRt*_1/2_ FDR < 0.1, or TR FDR > 0.45 and *TRt*_1/2_ FDR < 0.3.

## References

1 Hager, G. L., McNally, J. G. & Misteli, T. Transcription dynamics. Mol Cell 35, 741–753, doi:10.1016/j.molcel.2009.09.005 (2009).

2 Boettiger, A. N., Ralph, P. L. & Evans, S. N. Transcriptional regulation: effects of promoter proximal pausing on speed, synchrony and reliability. PLoS Comput Biol 7, e1001136, doi:10.1371/journal.pcbi.1001136 (2011).

3 Boettiger, A. N. Analytic approaches to stochastic gene expression in multicellular systems. Biophys J. 105, 2629–2640, doi:10.1016/j.bpj.2013.10.033 (2013).

4 Larson, D. R., Zenklusen, D., Wu, B., Chao, J. A. & Singer, R. H. Real-Time Observation of Transcription Initiation and Elongation on an Endogenous Yeast Gene. Science 332, 475–478 (2011).

5 Suter, D. M. et al. Mammalian Genes Are Transcribed with Widely Different Bursting Kinetics. Science 332, 472–474 (2011).

6 Karpova, T. S. et al. Concurrent Fast and Slow Cycling of a Transcriptional Activator at an Endogenous Promoter. Science 319, 466–469 (2008).

7 Poorey, K. et al. Measuring chromatin interaction dynamics on the second time scale at single-copy genes. Science 342, 369–372, doi:10.1126/science.1242369 (2013).

8 Viswanathan, R., Hoffman, E. A., Shetty, S. J., Bekiranov, S. & Auble, D. T. Analysis of chromatin binding dynamics using the crosslinking kinetics (CLK) method. Methods 70, 97–107, doi:10.1016/j.ymeth.2014.10.029 (2014).

9 Grimaldi, Y., Ferrari, P. & Strubin, M. Independent RNA polymerase II preinitiation complex dynamics and nucleosome turnover at promoter sites in vivo. Genome Res 24, 117–124, doi:10.1101/gr,157792.113 (2014).

10 Haruki, H., Nishikawa, J. & Laemmli, U. K. The anchor-away technique: rapid, conditional establishment of yeast mutant phenotypes. Mol Cell 31, 925–932, doi:10.1016/j.molcel.2008.07.020 (2008).

11 Chen, J. et al. Single-molecule dynamics of enhanceosome assembly in embryonic stem cells. Cell 156, 1274–1285, doi:10.1016/j.cell.2014.01.062 (2014).

12 Sprouse, R. O. et al. Regulation of TATA-binding protein dynamics in living yeast cells. Proc Natl Acad Sci USA 105, 13304–13308, doi:10.1073/pnas.0801901105 (2008).

13 Dion, M. F. et al. Dynamics of replication-independent histone turnover in budding yeast. Science 315, 1405–1408, doi:10.1126/science.1134053 (2007).

14 Lickwar, C. R., Mueller, F., Hanlon, S. E., McNally, J. G. & Lieb, J. D. Genome-wide protein-DNA binding dynamics suggest a molecular clutch for transcription factor function. Nature 484, 251–255, doi:10.1038/nature10985 (2012).

15 van Werven, F. J., van Teeffelen, H. A., Holstege, F. C. & Timmers, H. T. Distinct promoter dynamics of the basal transcription factor TBP across the yeast genome. Nat Struct Mol Biol 16, 1043–1048, doi:10.1038/nsmb.1674 (2009).

16 Grunberg, S. & Hahn, S. Structural insights into transcription initiation by RNA polymerase II. Trends Biochem Sci 38, 603–611, doi:10.1016/j.tibs.2013.09.002 (2013).

17 Pelechano, V., Chavez, S. & Perez-Ortin, J. E. A complete set of nascent transcription rates for yeast genes. PLoS One 5, e15442, doi:10.1371/journal.pone.0015442 (2010).

18 Rando, O. J. & Winston, F. Chromatin and transcription in yeast. Genetics 190, 351–387, doi:10.1534/genetics. 111.132266 (2012).

19 Nock, A., Ascano, J. M., Barrero, M. J. & Malik, S. Mediator-regulated transcription through the +1 nucleosome. Mol Cell 48, 837–848, doi:10.1016/j.molcel.2012.10.009 (2012).

20 Teves, S. S., Weber, C. M. & Henikoff, S. Transcribing through the nucleosome. Trends Biochem Sci 39, 577–586, doi:10.1016/j.tibs.2014.10.004 (2014).

21 Venters, B. J. et al. A comprehensive genomic binding map of gene and chromatin regulatory proteins in Saccharomyces. Mol Cell 41, 480–492, doi:10.1016/j.molcel.2011.01.015 (2011).

22 Basehoar, A. D., Zanton, S. J. & Pugh, B. F. Identification and distinct regulation of yeast TATA box-containing genes. Cell 116, 699–709 (2004).

23 Moqtaderi, Z. & Struhl, K. Genome-wide occupancy profile of the RNA polymerase III machinery in Saccharomyces cerevisiae reveals loci with incomplete transcription complexes. Mol Cell Biol 24, 4118–4127 (2004).

24 Stewart-Ornstein, J., Weissman, J. S. & El-Samad, H. Cellular noise regulons underlie fluctuations in Saccharomyces cerevisiae. Mol Cell 45, 483–493, doi:10.1016/j.molcel.2011.11.035 (2012).

25 Ravarani, C. N., Chalancon, G., Breker, M., de Groot, N. S. & Babu, M. M. Affinity and competition for TBP are molecular determinants of gene expression noise. NatCommun 7, 10417, doi:10.1038/ncomms10417 (2016).

26 Yudkovsky, N., Ranish, J. A. & Hahn, S. A transcription reinitiation intermediate that is stabilized by activator. Nature 408, 225–229, doi:10.1038/35041603 (2000).

27 Sainsbury, S., Bernecky, C. & Cramer, P. Structural basis of transcription initiation by RNA polymerase II. Nat Rev Mol Cell Biol 16, 129–143, doi:10.1038/nrm3952 (2015).

28 Santisteban, M. S., Hang, M. & Smith, M. M. Histone variant H2A.Z and RNA polymerase II transcription elongation. Mol Cell Biol 31, 1848–1860, doi:10.1128/MCB.01346-10 (2011).

29 Mencia, M., Moqtaderi, Z., Geisberg, J. V., Kuras, L. & Struhl, K. Activator-specific recruitment of TFIID and regulation of ribosomal protein genes in yeast. Mol Cell 9, 823–833 (2002).

30 Reiner, A., Yekutieli, D. & Benjamini, Y. Identifying differentially expressed genes using false discovery rate controlling procedures. Bioinformatics 19, 368–375 (2003).

